# Glioma-derived CCL2 and CCL7 mediate migration of immune suppressive CCR2^+^ myeloid cells into the tumor microenvironment in a redundant manner

**DOI:** 10.1101/2022.07.08.499173

**Authors:** Gregory P. Takacs, Christian J. Kreiger, Defang Luo, Guimei Tian, Julia S. Garcia, Loic P. Deleyrolle, Duane A. Mitchell, Jeffrey K. Harrison

**Affiliations:** Department of Pharmacology & Therapeutics, University of Florida College of Medicine, Gainesville, FL, USA; Department of Neurosurgery, University of Florida College of Medicine, Gainesville, FL, USA

**Keywords:** Glioma, Chemokine, Chemokine Receptor, Migration, Immune-suppression, Myeloid, MDSC, Bone Marrow

## Abstract

Glioblastoma (GBM) is the most common and malignant primary brain tumor, resulting in poor survival despite aggressive therapies. GBM is characterized in part by a highly heterogeneous and immunosuppressive tumor microenvironment (TME) made up predominantly of infiltrating peripheral immune cells. One significant immune cell type that contributes to glioma immune evasion is a population of immunosuppressive, hematopoietic cells, termed myeloid-derived suppressor cells (MDSCs). Previous studies suggest that a potent subset of myeloid cells, expressing monocytic (M)-MDSC markers, distinguished by dual expression of chemokine receptors CCR2 and CX3CR1, utilize CCR2 to infiltrate into the TME. This study evaluated the T cell suppressive function and migratory properties of CCR2^+^/CX3CR1^+^ MDSCs. Bone marrow-derived CCR2^+^/CX3CR1^+^ cells adopt an immune suppressive cell phenotype when cultured with glioma-derived factors. Recombinant and glioma-derived CCL2 and CCL7 induce the migration of CCR2^+^/CX3CR1^+^ MDSCs with similar efficacy. KR158B-CCL2 and -CCL7 knockdown murine gliomas contain equivalent percentages of CCR2^+^/CX3CR1^+^ MDSCs compared to KR158B gliomas. Combined neutralization of CCL2 and CCL7 completely blocks CCR2-expressing cell migration to KR158B cell conditioned media. High levels of CCL2 and CCL7 are also associated with negative prognostic outcomes in GBM patients. These data provide a more comprehensive understanding of the function of CCR2^+^/CX3CR1^+^ MDSCs and the role of CCL2 and CCL7 in the recruitment of these immune suppressive cells and further support the significance of targeting this chemokine axis in GBM.

## Introduction

Glioblastoma (GBM) is a highly aggressive and recurrent primary brain tumor that continues to challenge patients and oncologists as current interventions are minimally effective (1,2). Currently, standard of care therapy relies on surgical resection of the tumor mass followed by focal radiation and chemotherapy (temozolomide) (3–5). Foremost in GBM patients, the immune suppressive tumor microenvironment contributes to immune evasion, disease progression, and poor overall survival (6– 9). Attempts at harnessing anti-tumor immune responses to overcome the immunosuppressive microenvironment have been made in cancer therapy (10–12). For example, clinically successful immunotherapy has targeted immune checkpoint systems, including the programmed cell death protein-1, i.e., PD-L1/PD-1, pathway. Unfortunately, to date targeting the PD-L1/PD-1 axis in human gliomas has not demonstrated efficacy as an adjuvant monotherapy (13,14). While the mechanism by which gliomas are resistant to PD-1 blockade is not entirely resolved, mounting evidence suggests that infiltrating immune suppressive cells contribute significantly to the resistant phenotype (15–17).

Chief amongst the immune suppressive cells which gain access to the glioma microenvironment are a subset of myeloid cells termed myeloid-derived suppressor cells (MDSCs) (15). In humans, MDSCs represent a heterogeneous cell population that are delineated into three major classes based on phenotypic and morphological features: early-stage (e), polymorphonuclear (PMN/G)-, and monocytic (M)-MDSCs (18,19). Although M-MDSCs are present within the human glioma microenvironment in a lower proportion than are the other subtypes, M-MDSCs display the most potent immune suppressive phenotype through deleterious anti-tumor lymphocyte interactions. M-MDSCs suppress lymphocytes via production of free radicals and enzymes that deplete essential lymphocyte metabolites (20–23). In murine gliomas, M-MDSCs are the predominant subtype and are characterized by lineage markers CD45^+^, CD11b^+^, Ly6C^hi^, and Ly6G^−^ (19,24,25).

We have previously reported that three populations of myeloid cells are identified in the glioma microenvironment according to their expression of chemokine receptors CCR2 and CX3CR1. One of these populations, co-expressing chemokine receptors CCR2 and CX3CR1 (denoted as CCR2^+^/CX3CR1^+^), express markers consistent with M-MDSCs. Pharmacologic or genetic targeting of CCR2-expressing cells via a CCR2 antagonist or gene deletion limited the presence of these cells within the tumor and promoted their sequestration within the bone marrow. In combination with the immune checkpoint inhibitor, αPD-1, CCR2 antagonism unmasked an effect of PD-1 blockade in slowing the tumor progression of two immune checkpoint inhibitor-resistant murine gliomas (KR158B and 005 GSC) (26). While these previous findings established that CCR2^+^/CX3CR1^+^ MDSCs utilize CCR2 to traffic into the glioma microenvironment, it is unclear what chemokines drive this CCR2-dependent migration. This study investigated the T cell suppressive function and chemokine ligand dependency by which CCR2^+^/CX3CR1^+^ M-MDSCs traffic into the glioma microenvironment. Using a preclinical glioma model, we demonstrate that CCR2^+^/CX3CR1^+^ cells are sourced from the bone marrow, suppress both CD4^+^ and CD8^+^ T cells, and migrate to CCL2 and/or CCL7 in a CCR2-dependent manner. We also identify CCL2 and CCL7 as predictors of survival in human glioblastoma. These data establish the immune suppressive and migratory properties of CCR2^+^/CX3CR1^+^ myeloid cells and confirm their role as glioma-associated M-MDSCs.

## Methods

### Animals

*Ccr2*^*RFP/WT*^*/Cx3cr1*^*GFP/WT*^ mice were generated through the breeding of Ccr2-deficient (*Ccr2*^*RFP/RFP*^*[B6*.*129(Cg)-Ccr2*^*tm2*.*1Ifc*^*/J*]), and Cx3cr1-deficient (*Cx3cr1*^*GFP/GFP*^*[B6*.*129P-Cx3cr1*^*tm1Litt*^*/J*]) mice. Wildtype C57BL/6, Ccr2-deficient, and Cx3cr1-deficient mice were purchased from The Jackson Laboratory. All procedures involving animal housing and surgical protocols were followed according to the guidelines of the University of Florida Institutional Animal Care and Use Committee.

### Generation of Chimeric Mice

Chimeric mice were generated through a bone marrow transplant of *Ccr2*^*WT/RFP*^*/Cx3cr1*^*WT/GFP*^ donor mice into wildtype C57BL/6 recipient mice. Wildtype mice were placed under anesthesia (Xylazine 0.5mL, Ketamine 0.7mL, Saline 5.6mL) through intra peritoneal injection (100μL/20g mouse). Subsequently, wildtype mice received 900 cGy x-ray radiation (X-RAD 350 irradiator). Bone marrow was prepared from *Ccr2*^*WT/RFP*^*/Cx3cr1*^*WT/GFP*^ mice as described below. Cells were diluted to a final concentration of 10,000 cells/μL. After irradiation (∼4hrs), whole bone marrow from *Ccr2*^*WT/RFP*^*/Cx3cr1*^*WT/GFP*^ donor mice was tail vein injected (100 μL) into irradiated wildtype C57BL/6 recipient mice. Seven days post-irradiation, Baytril (fluoroquinolone antibiotic) was added to the drinking water at 0.5 mg/ml for two weeks. Following recovery, chimeric mice were implanted with KR158B gliomas (see “Orthotopic Brain Tumor Model”) and evaluated via flow cytometry (see “Flow Cytometry Analysis”)

### Orthotopic Brain Tumor Model

Animals were anesthetized using isoflurane and administered analgesia prior to cell injection. While under anesthesia, the surgical site was prepared and a 2-to 3-mm incision was made at the midline of the skull. Using a stereotaxic apparatus (Stoelting), the mice were secured, and a Hamilton syringe was positioned 2-mm lateral from the bregma. KR158B, KR158B CCL2 knockdown, or KR158B CCL7 knockdown glioma cells (3.5 × 10^4^ in a total volume of 2 μL) were injected 3-mm deep into the right cerebral hemisphere using an automated microfluidic injection system (Stoelting) at a rate of 1 μL /min; cells were suspended in a 1:1 ratio of methylcellulose to PBS. Post-injection, the needle was retracted slowly, and the surgical site was closed via suture and bone wax. Animals were then placed into a warm cage for postsurgical monitoring.

### Tissue Isolation

Mice were euthanized at experimental endpoint. Transcardial perfusions, using a 10mL syringe with a 25G winged infusion set, of 20mL 0.9% saline solution were administered to remove intra-vasculature associated cells. Femurs, tibiae, and humeri were harvested from the animal. Fat and muscle were removed, and the bones were subsequently cut at one end to expose bone marrow. Bones were placed in microcentrifuge tubes (2 bones per tube) with the bottoms pierced and nested in 1.5mL centrifuge tubes containing 100uL PBS. Tubes were centrifuged at 5,700 x g for 20 seconds to flush the bone marrow. Spleens were harvested and placed on a petri dish. Fat was trimmed from the tissue and spleens were injected with 1mL PBS via 18G needle. Spleens were minced using a razor blade and transferred to 15mL conical tubes containing 5mL PBS. Using a 5mL syringe and 18G needle, the tissue was mechanically dissociated via passage through the needle 20 times. Splenocytes were collected via centrifugation (4 °C, 380 × *g*, 5 min). Bone marrow cells and splenocytes were resuspended in 1mL Ammonium-Chloride-Potassium (ACK) Lysis buffer (Gibco, Invitrogen) and placed on ice for 5 min to lyse red blood cells. Subsequently, lysis buffer was quenched with 5mL fluorescence-activated cell-sorter (FACS) washing buffer (1% FBS in PBS) and strained through a 40-μm cell strainer. Cells were collected via centrifugation (4 °C, 380 × *g*, 5 min) and counted by trypan blue exclusion. Brains were removed and tumors were extracted and mechanically minced using a razor blade. Tumors were placed in 4 °C Accumax dissociation solution (Innovative Cell Technologies) and incubated at 37 °C for 5 min, followed by 5 min of agitation at room temperature. Cells were then passed through a 40-μm strainer, centrifuged (4 °C, 380 × *g*, 5 min), and resuspended in 4 mL of 70% Percoll (70% Percoll and 1% PBS in RPMI-1640 cell medium). The 70% Percoll/cell solution was then carefully layered beneath 37% Percoll layer (4 mL, 37% Percoll and 1% PBS in RPMI-1640 cell medium) using an 18-gauge needle. Samples were then centrifuged for 30min at room temperature (500 x g). Cells at the interface were collected and transferred into a 1.5 mL microcentrifuge tube. Cells were washed with cold PBS and counted by trypan blue exclusion.

### Flow Cytometry Analysis

Single cell suspensions were prepared from tissues as described above and diluted to 1 × 10^6^ cells/100uL. Subsequently, cells were stained for markers of interest (see Supplement Table 1) for 30 min at 4 °C. Cells were then washed twice in ice-cold PBS and stained with a viability dye. Stained samples were analyzed using single-color compensation on a Sony SP6800 spectral analyzer or Beckman Coulter CytoFLEX LX 96-well plate system and quantified using FlowJo V10.8.1 (BD Biosciences).

### Cell Culture

KR158B, KR158B-Luciferase, KR158B CCL2 knockdown, and KR158B CCL7 knockdown glioma cells were cultured in Dulbecco’s Modified Eagle Medium (DMEM) supplemented with 1% penicillin-streptomycin and 10% fetal bovine serum (FBS). Cells were grown in a humidified incubator at 37 °C with 5% CO_2_. DMEM and penicillin-streptomycin were purchased through Invitrogen. FBS was purchased through Thermo Scientific.

### Generation of CCL2- and CCL7-Deficient Glioma Cells

Plasmids for knockdown of CCL2 (TRCN0000301701 and TRCN0000301702) and CCL7 (TRCN0000317599, TRCN0000068135, and TRCN0000068136) were obtained from Sigma. shRNA control plasmid (SHC002. Sigma) was used as non-targeting control. ShRNA plasmids were purified with QIAprep Spin Miniprep Kit (#27106, Qiagen) after overnight incubation with E-coli bacteria. Packaging 293T/17 cells were co-transfected with the different shRNAs and the packaging plasmids psPAX2 and pMD2.G, to generate viral particles, which were subsequently used to transduce KR158B cells. KR158B CCL2 knockdown were generated using the combination of TRCN0000301701 and TRCN0000301702. TRCN0000317599, TRCN0000068135, and TRCN0000068136 were combined to generate KR158B CCL7 knockdown.

### Cytokine Quantification

Cytokine quantification was completed using mouse CCL2 (Invitrogen# 88-7391-22) and mouse CCL7 (Invitrogen# BMS6006INST) enzyme-linked immunosorbent assay (ELISA) analysis following manufacturer protocols. KR158B, KR158B CCL2 knockdown and KR158B CCL7 knockdown glioma cell lines were cultured to 90% confluency. Cells were counted and plated in a 96-well plate at 50, 100 or 500 cells/uL in 200uL complete DMEM media. Cells were incubated at 37°C, 5% CO_2_ for 24 hours. Following incubation, well contents were transferred to tubes and centrifuged as previously described. Supernatant was aliquoted and frozen at -80°C until use for ELISA as previously described.

### Mouse Brain Fixation and Immunohistochemistry

Transcardial perfusions, using a 10mL syringe with a 25G winged infusion set, of 20mL 4.0% paraformaldehyde (PFA) solution were administered. Following fixative perfusion, mouse brains were removed and soaked in 4.0% PFA for 1hr. Brains were subsequently transferred to 30% sucrose solution for 24hrs and snap frozen using liquid nitrogen chilled 2-Methylbutane. Brains were embedded in optimal cutting temperature compound and mounted for cryo-sectioning (Lecia Biosystems Cryostat). 5-10μm thick sections were taken and mounted on microscope slides. Sections were dried overnight at 4°C. Tissue sections were brought to room-temperature and washed 3 times in PBS and counterstained with antifade mounting medium with DAPI (Vectashield). Brain tumor sections were imaged using an inverted Nikon TiE-PFS-A1R confocal microscope. Images were post-processed using Nikon Elements software.

### Bone Marrow Culture

Induction of MDSCs was adapted from previously published work (Alban et al.) (27). Bone marrow-derived cells from wildtype C57BL/6 mice were prepared as previously described. Cells were then plated at a density of 400,000 cells/cm^2^ and concentration of 1,000 cells/uL in media consisting of 50% complete RPMI (RPMI + 10% FBS + 2mM L-Glutamine) and 50% KR158B conditioned media. Additionally, the media was supplemented with 40ng/mL GM-CSF (R&D 415-ML) and 40ng/mL IL-6 (R&D 406-ML). On day 5, suspended cells were collected, the flask was washed in PBS and scraped using a cell scraper (Fisher), and all contents were joined together in a 50mL conical tube. Cells were collected via centrifugation (4 °C, 380 × *g*, 5 min) and counted by trypan blue exclusion. Cells were then either subjected to flow cytometry (see “Flow Cytometry Analysis”) or utilized for the T cell suppression assay (see “T cell Suppression Assay”).

### T cell Suppression Assay

Following a 5-day culture (see “Bone Marrow Culture”), MDSC enriched bone marrow cells were collected from culture as described above and subjected to M-MDSC magnetic bead isolation (Miltenyi Biotec) according to manufacturer’s protocols. Additionally, fresh splenocytes were isolated as previously described and subjected to Pan-T cell magnetic bead isolation (Miltenyi Biotec) according to manufacturer’s protocols. Following isolation, T cells were collected via centrifugation and resuspended at a density of 1 million cells/mL in PBS. T cells were incubated with 1uL CellTrace FarRed Cell Proliferation dye (ThermoFisher C34564) per 1 million cells for 20 minutes at RT. Following incubation, the dye was quenched in 5 times the present volume of complete RPMI. T cells were collected via centrifugation (4 °C, 380 × *g*, 5 min) and resuspended in complete RPMI at 1,000 cells/uL. Dynabeads Mouse T Activator CD3/CD28 beads (Thermofisher 11452D) were washed in complete RPMI and mixed with stained T cells at a 2:1 (activating bead:T cell) ratio. T cells were retained at each step to ensure for unstained, unstimulated, and stained/unstimulated controls. 100,000 T cells were added per well in a round-bottom 96-well plate and MDSCs were added at ratios of 1:4, 1:2 and 1:1 (MDSCs:T cells). Co-cultures were incubated at 37°C for 3 days. Following incubation, well contents, in addition to 2 subsequent PBS well washes, were transferred to centrifuge tubes. Tubes were then placed on the Dynamag-2 (Thermofisher 12321D) to remove activating beads. Cells were collected by centrifugation and stained for CD3, CD4 and CD8 (Biolegend 100234; 100510; 100708) for flow cytometry analysis. Each biologic and condition were run in triplicate. Technical triplicates were averaged prior to statistical analysis.

### In Vitro Cell Migration

Bone marrow cells were isolated from *Ccr2*^*WT/RFP*^*/Cx3cr1*^*WT/GFP*^ mice as described previously. Cells were diluted to a final concentration of 2,000 cells/μL in migration buffer consisting of RPMI-1640, 25mM HEPES, 1% penicillin-streptomycin, and 0.1% BSA (>98% quality). In vitro migration of CCR2^WT/RFP^/CX3CR1^WT/GFP^ cells was assessed using a transwell-96 well plate with 5μm polycarbonate membrane (Corning; product number 3388). Recombinant Mouse CCL2, CCL7, and soluble CX3CL1 chemokines were purchased from R & D Systems (product numbers 479-JE-010; 456-MC-010; 571-MF-025). Recombinant proteins were reconstituted following manufacture preparation and storage guidelines. Recombinant CCL2, CCL7, and soluble CX3CL1 ligands were diluted in migration buffer and seeded at 150μL/well in the bottom chamber. To validate a chemotaxis effect, chemokine was also placed in the top and bottom chambers at equivalent concentrations (i.e., 30ng/mL top chamber and 30ng/mL bottom chamber). Cells were plated at 150μL/well in the top chamber, and the plate was incubated at 37°C, 5% CO_2_ for 2 hours. After incubation, the membrane insert was lightly shaken to detach migrated cells on the underside of the membrane and then discarded. Wells were analyzed for CCR2^WT/RFP^ and CCR2^WT/RFP^/CX3CR1^WT/GFP^ populations using single color compensation on Beckman Coulter CytoFLEX LX 96-well plate system. 75uL/well was collected at a flow rate of 150uL/min with 3s shake time and backflush between wells. Gating strategy proceeded as follows: 1. Positive gate for myeloid population according to forward-scatter area (FSC-A) and side-scatter area (SSC-A). 2. Doublet exclusion according to FSC-A and forward-scatter height (FSC-H). 3. CCR2/CX3CR1 co-expression according to PE and FITC channels. Gating strategy was established according to analysis of raw bone marrow samples and applied constantly throughout the analysis. Final gating analysis was conducted using FCS Express software (De Novo) or FlowJo V10.8.1 (BD Biosciences). Control wells containing no chemokine in the bottom well were averaged and normalized as 100% migration. Sample wells are compared relative to control wells for presentation and statistical analyses.

For analysis of migration to conditioned media, KR158B or KR158B CCL2 KD cells were cultured to 90% confluency in a tissue-culture T-75 flask containing complete DMEM (see “Cell Culture” section in methods). Media was washed out and cells were plated in a 6-well plate in 6mL migration buffer at 50, 100, and 500 cells/μL overnight at 37 °C, 5% CO_2_. Contents from wells were extracted and centrifuged at 1,000 RPM for 5 minutes. Supernatant was collected, filtered, and aliquoted in the bottom chamber of the transwell-96 well plate. For neutralization experiments, polyclonal goat IgG antibodies (anti-CCL2, anti-CCL7, and normal goat IgG control) were purchased from R & D Systems (product number AB-479-NA, AF-456-NA, and AB-108-C) and stored following manufacture instructions. All wells that received neutralizing antibodies received equal quantities of exogenous protein (i.e., 8.25 μg/well) through supplementation of normal goat IgG control. Raw bone marrow was seeded in the top chamber as previously mentioned. Each biologic and condition were run in triplicate. Technical triplicates were averaged prior to statistical analysis.

### Survival Analysis

The complete human glioblastoma multiforme (GBM) patient dataset was mined from The Cancer Genome Atlas (TCGA Research Network: cancer.gov/tcga) The Georgetown Database of Cancer (G-DOC) platform to extract gene expression and clinical parameters (28–30). G-DOC platform was accessed on February 4, 2022. Gene expression was gathered from the Affymetrix dataset (Affymetrix HT Human Genome U133a microarray platform by the Broad Institute of MIT and Harvard University cancer genomic characterization center) and RNA sequencing dataset (Illumina HiSeq 2000 RNA Sequencing platform by the University of North Carolina TCGA genome characterization center). Patients were stratified into low or high CCL2, CCL7, and CCL2 ∩ CCL7 expressing categories (LOW<25^th^ percentile and HIGH>75^th^ percentile, respectively). Percentiles were generated using descriptive statistics function in GraphPad Prism version 9.3.1 for Windows, GraphPad Software, San Diego, California USA. Survival curve comparisons and numbers at risk were calculated using Log-rank (Mantel-Cox test) and graphically illustrated through GraphPad Prism version 9.3.1. P-values are reported in figures.

### Statistical Analysis

Multiple t-tests, Log-rank (Mantel-Cox), one-way ANOVA and two-way ANOVA analyses were performed in GraphPad Prism version 9.3.1 to determine statistically significant differences between groups. Multiple comparisons were corrected for with the recommended Dunnett multiple comparison test. A p-value <0.05 was considered significant and is indicated by symbols depicted in the figures, figure legends and text.

## Results

### CCR2^+^/CX3CR1^+^ cells in the glioma microenvironment are sourced from the bone marrow

We previously established that a glioma-associated CCR2^+^/CX3CR1^+^ myeloid cell population also expresses markers consistent with M-MDSCs. A CCR2^+^/CX3CR1^+^ myeloid cell population, expressing the same MDSC markers, is also present in bone marrow. To examine the T cell suppressive and migratory properties of these CCR2^+^/CX3CR1^+^ cells, dual transgenic *Ccr2*^*WT/RFP*^*/Cx3cr1*^*WT/GFP*^ mice were utilized in order to facilitate the direct examination of CCR2- and CX3CR1-expressing cells. Fluorescent confocal microscopy of intracranial KR158B tumors confirmed the presence of brain-resident CX3CR1^WT/GFP^ microglia and revealed that CCR2^WT/RFP^ and CCR2^WT/RFP^/CX3CR1^WT/GFP^ cells were also present within the TME as early as 5 days post-implantation of KR158B tumor cells (**Figure 1A**). Fluorescent imaging of naïve (non-tumor) brain tissue confirmed the absence of any RFP-expressing cells in non-tumor bearing brain tissue while RFP/GFP positive cells were present in bone marrow (**Figure 1A**).

**Figure 1.**
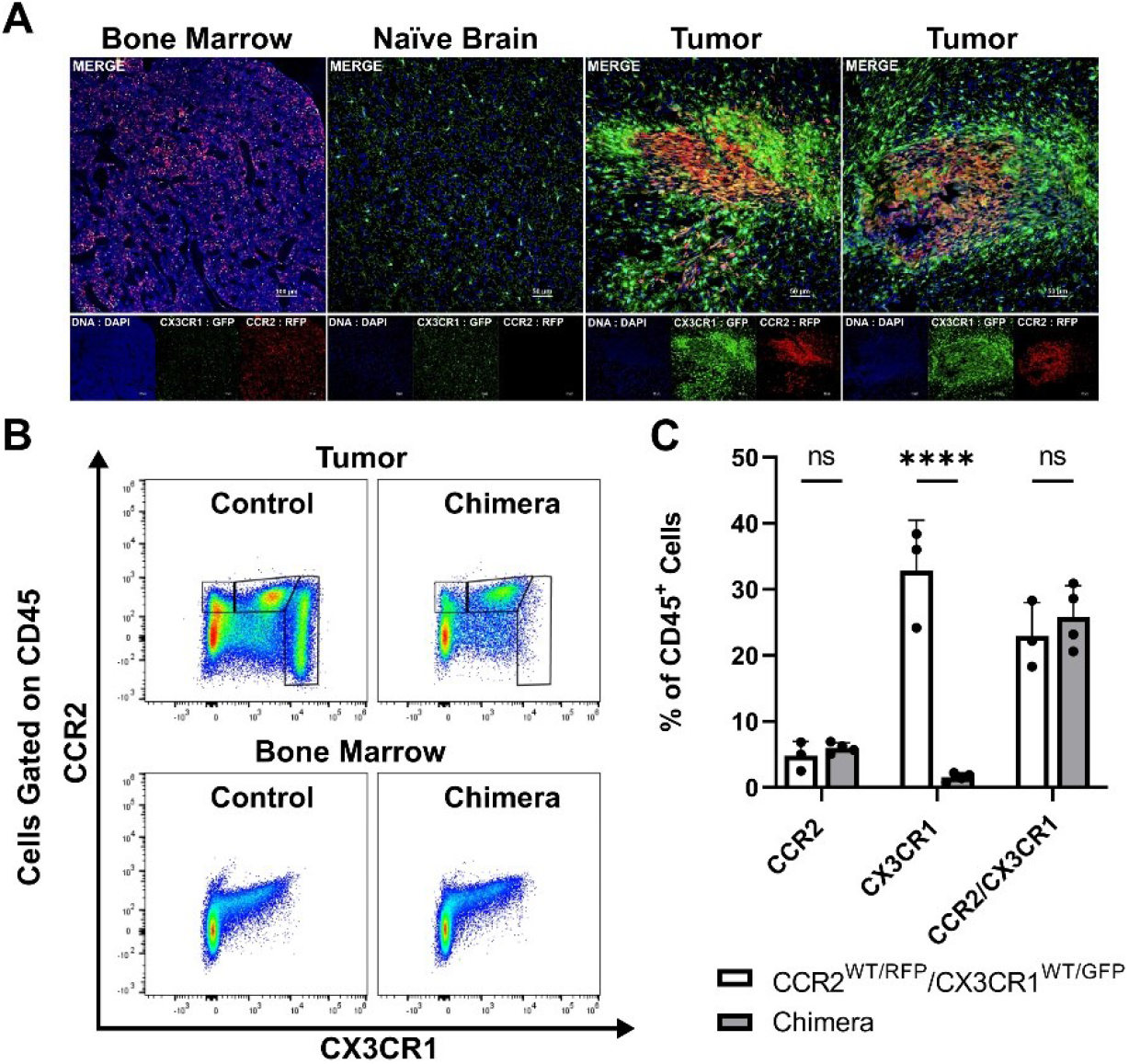
CCR2^+^/CX3CR1^+^ cells infiltrate the glioma microenvironment and are derived from the bone marrow. (**A)** Representative immunofluorescent images of bone marrow, naïve brain, and tumor-implanted brains 5 days post-implantation. Images depict the absence of CCR2^WT/RFP^ cells within normal brain and presence within bone marrow and tumors 5 days post-implantation. (**B)** Representative flow cytometry plots of tumors and bone-marrow of non-irradiated control and chimeric animals at experimental endpoint. **(C)** Quantification of CCR2^+^, CX3CR1^+^ and CCR2^+^/CX3CR1^+^ leukocytes in control and chimeric animals (n=3). GraphPad Prism was used to conduct two-way ANOVA statistics (Dunnett’s multiple comparisons test). Differences are compared to the control (0) condition. p-values: 0.0332(*), 0.0021(**), 0.0002(***), <0.0001(****)

To directly establish if CCR2^+^/CX3CR1^+^ cells present within KR158B gliomas are sourced from the bone marrow we generated chimeric mice harboring *Ccr2*^*WT/RFP*^*/Cx3cr1*^*WT/GFP*^ bone marrow cells. Irradiated wildtype C57BL/6 mice (recipient) received whole bone marrow isolated from *Ccr2*^*WT/RFP*^*/Cx3cr1*^*WT/GFP*^ mice (donor) and, following immune reconstitution, chimeric mice were orthotopically implanted with KR158B glioma cells. At experimental endpoint, bone marrow and brain tumor tissue were processed for flow cytometry. Flow cytometry analysis identified the presence of CCR2^+^ and CCR2^+^/CX3CR1^+^ cells in the tumors of chimeric mice (p<.0001) (**Figure 1B, C**) which indicates that these populations were derived from the bone marrow. GFP+ cells were absent from these tumors, suggesting that this population is brain-derived.

### CCR2^+^/CX3CR1^+^ cells suppress CD8^+^ and CD4^+^ T cell proliferation and IFN-γ production

To investigate the functionality of CCR2^+^/CX3CR1^+^ cells, the impact on T cell proliferation and function was assessed. Having determined that CCR2^+^/CX3CR1^+^ cells are bone marrow-derived, whole bone marrow was harvested and cells were cultured in the presence of KR158B glioma-derived factors (conditioned media) containing soluble GM-CSF and IL-6 to enrich and expand the population of dual-expressing chemokine receptor cells. Following magnetic bead MDSC isolation, flow cytometry analysis confirmed the isolation of cells expressing CD45, CD11b, Ly6C, and chemokine receptors CCR2 and CX3CR1 (**Figure 2A**); cells were negative for Ly6G. The enriched, bone marrow-derived cells significantly suppressed the proliferation of both CD4^+^ and CD8^+^ T cells at ratios 1:2 and 1:1 (**Figure 2B-D**). In the presence of MDSCs at a 1:2 ratio, CD4^+^ T cell proliferation decreased from 71% to 39% while CD8^+^ T cell proliferation decreased from 82% to 50%. When co-cultured at a 1:1 ratio of MDSCs to T cells, CD4^+^ and CD8^+^ T cell proliferation was suppressed to 20% and 18%, respectively (**Figure 2C, D)**. To further assess suppression within the co-culture, we analyzed the media from the suppression assay for levels of IFN-γ. Consistent with the results for proliferation, higher ratios of MDSCs:T cells also yielded lower concentrations of IFN-γ within the co-culture (**Figure 2E**), suggesting functional inhibition of effector T cells by the CCR2^+^/CX3CR1^+^ MDSCs. These results establish that bone marrow CCR2^+^/CX3CR1^+^ cells, when incubated in glioma-derived factors, acquire a phenotype capable of disrupting the proliferation and function of both CD4- and CD8-expressing T cells.

**Figure 2.**
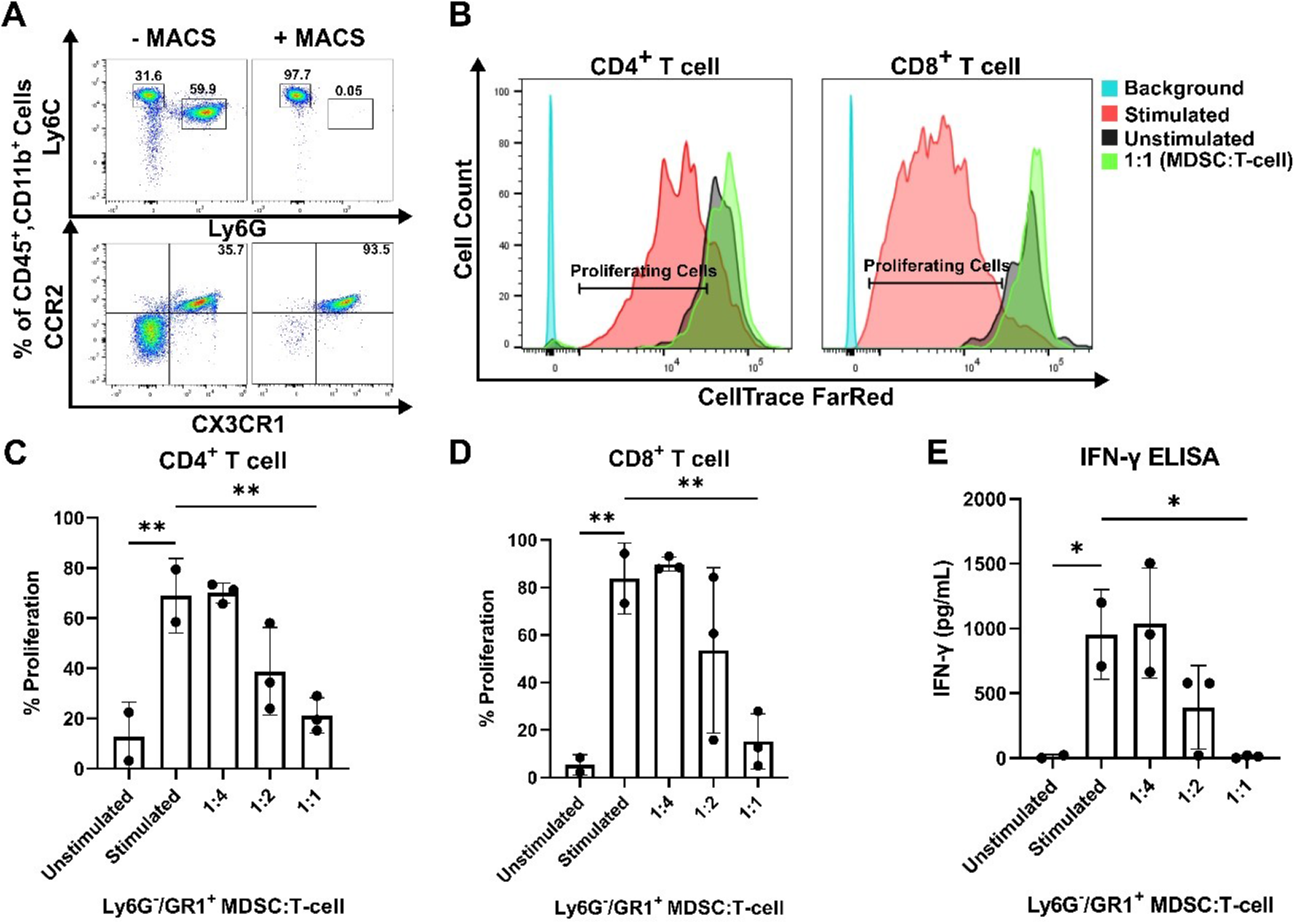
CCR2^+^/CX3CR1^+^ cells suppress CD8 and CD4 T cell proliferation and IFN-γ production. **(A)** Representative flow cytometry plot establishing CCR2^+^/CX3CR1^+^ cells are enriched using Ly6G and GR-1 magnetic-activated cell sorting (MACS). **(B)** Representative flow cytometry plot denoting CD4^+^ and CD8^+^ proliferating cells in the presence or absence of CCR2^+^/CX3CR1^+^ enriched cells. Quantification of **(C)** CD4^+^ or **(D)** CD8^+^ T cell proliferation in the presence and absence of CD3/CD28 activation beads (stimulated/unstimulated) and stimulated T cells co-cultured with enriched CCR2^+^/CX3CR1^+^ cells. Enriched CCR2^+^/CX3CR1^+^ cells were plated at varying ratios to dye-loaded T cells (T cell numbers were held constant). After 3 days, proliferation was assessed using flow cytometry (n=3). **(E)** Supernatant from co-culture T cell suppression assay was collected and analyzed for IFN-γ protein via ELISA (n=3). One-way ANOVA statistical analysis was conducted (Dunnett’s multiple comparisons test). Differences are compared to the stimulated control condition. p-values: 0.0332(*), 0.0021(**), 0.0002(***), <0.0001(****)

### High CCL2 and CCL7 expression is associated with lower overall survival in human glioblastoma

Of the five known human ligands of CCR2, three are shared with mice, namely CCL2, CCL7 and CCL8. To determine the impact of these chemokines on the clinical prognosis of human GBM, gene expression and survival data from The Cancer Genome Atlas (TCGA) GBM cohort was analyzed. Patients were stratified into “Low” and “High” expressing categories based on the lowest and highest quartiles of expression. Kaplan-Meyer survival curves, derived from Affymetrix and IlluminaHighseq datasets, were generated and Log-rank tests were utilized to compare the survival distributions. Similar to findings of Chang et al., a statistically significant decrease in survival among patients with high expression of CCL2, compared to low-expressing patients, was evident in the Affymetrix gene expression dataset (p<0.0005) (**Figure 3A**) (31). Similar results based on CCL7 expression were identified (p=0.0417) (**Figure 3B**). Upon grouping cohorts of high expression of CCL2 and CCL7 and low expression of CCL2 and CCL7 (denoted as the intersection sign “∩”), there was a statistically significant decrease in survival among the high expression cohort (p=0.0255) (**Figure 3C**). More striking results were revealed when analyzing the Illumina Highseq dataset. High CCL2 expression (p=0.0109) (**Figure 3D**) and high CCL7 (p=0.0319) (**Figure 3E**) expression among patients correlated with a statistically significant reduction in survival. This survival disadvantage was most pronounced among patients with high expression of both CCL2 and CCL7 (p=0.0018) (**Figure 3F**). These results indicate that high expression of CCL2 and CCL7 is negatively correlated with survival in the context of human GBM. Expression of CCL8, the other shared CCR2 chemokine across species, was not associated with a significant survival disadvantage (**Supplementary Figure 3C**). CCL13 and CCL16, CCR2 ligands found only in humans, also did not show a significant survival disadvantage (**Supplementary Figure 3**). Taken together with our previous data, we posit that this significant difference in survival between low and high expressors is due in part to an elevated level of recruitment of immunosuppressive CCR2^+^/CX3CR1^+^ cells into the glioma microenvironment.

**Figure 3.**
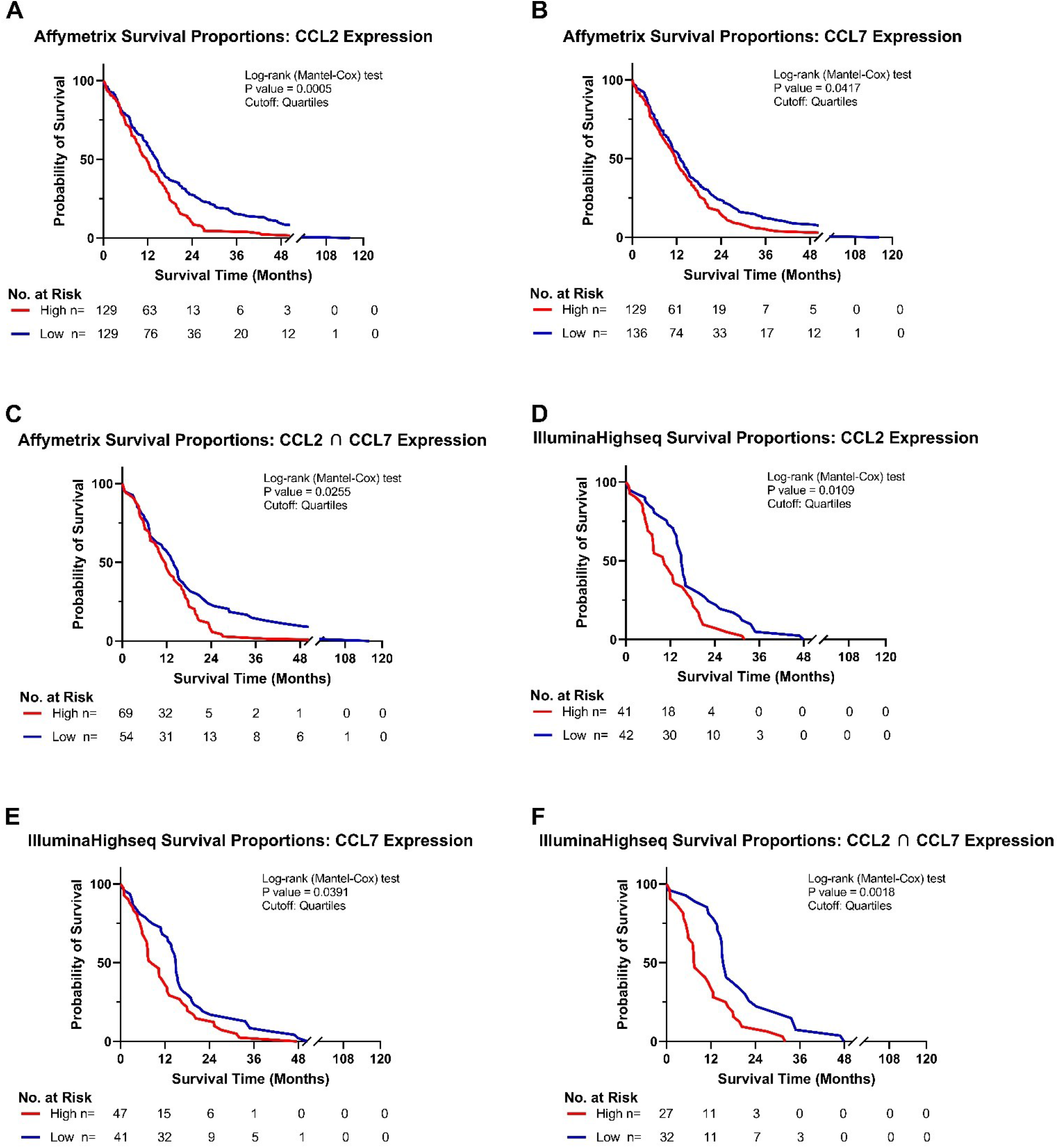
High CCL2 and CCL7 expression is associated with negative prognosis for patients with glioblastoma. **(A-C)** Kaplan-Meyer survival curves of GBM patients based on Affymetrix gene expression profiles of **(A)** CCL2 **(B)** CCL7 **(C)** intersection of CCL2 and CCL7 from TCGA database. **(D-F)** Kaplan-Meyer survival curves of GBM patients based on Illumina Highseq expression profiles of **(D)** CCL2 **(E)** CCL7 **(F)** intersection of CCL2 and CCL7 mined from TCGA database. High and low cohorts are stratified as top and bottom quartiles, respectively. Number at risk indicates surviving patients in each cohort at the respective timepoints of analysis. Log-rank (Mantel-Cox) test was conducted on high vs low expressing cohorts.

### CCR2^+^/CX3CR1^+^ cells migrate to recombinant CCL2 and CCL7 through CCR2

With evidence that CCR2^+^/CX3CR1^+^ cells represent a potent T cell suppressive population and CCR2 ligands (CCL2 and CCL7) confer poor survival in human GBM, we evaluated the impact of CCL2 and CCL7 on cell migration. To determine the migratory capacity of the CCR2^+^/CX3CR1^+^ cell population to chemokine ligands, a 96-well 5μm transwell migration assay was employed. Migration of CCR2^+^/CX3CR1^+^ cells was determined using flow cytometry, gating for CCR2^WT/RFP^ and CX3CR1^WT/GFP^ double-positive cells (**Figure 4A**) with results presented as percent migration relative to the control condition i.e., no recombinant chemokine in the bottom chamber.

**Figure 4.**
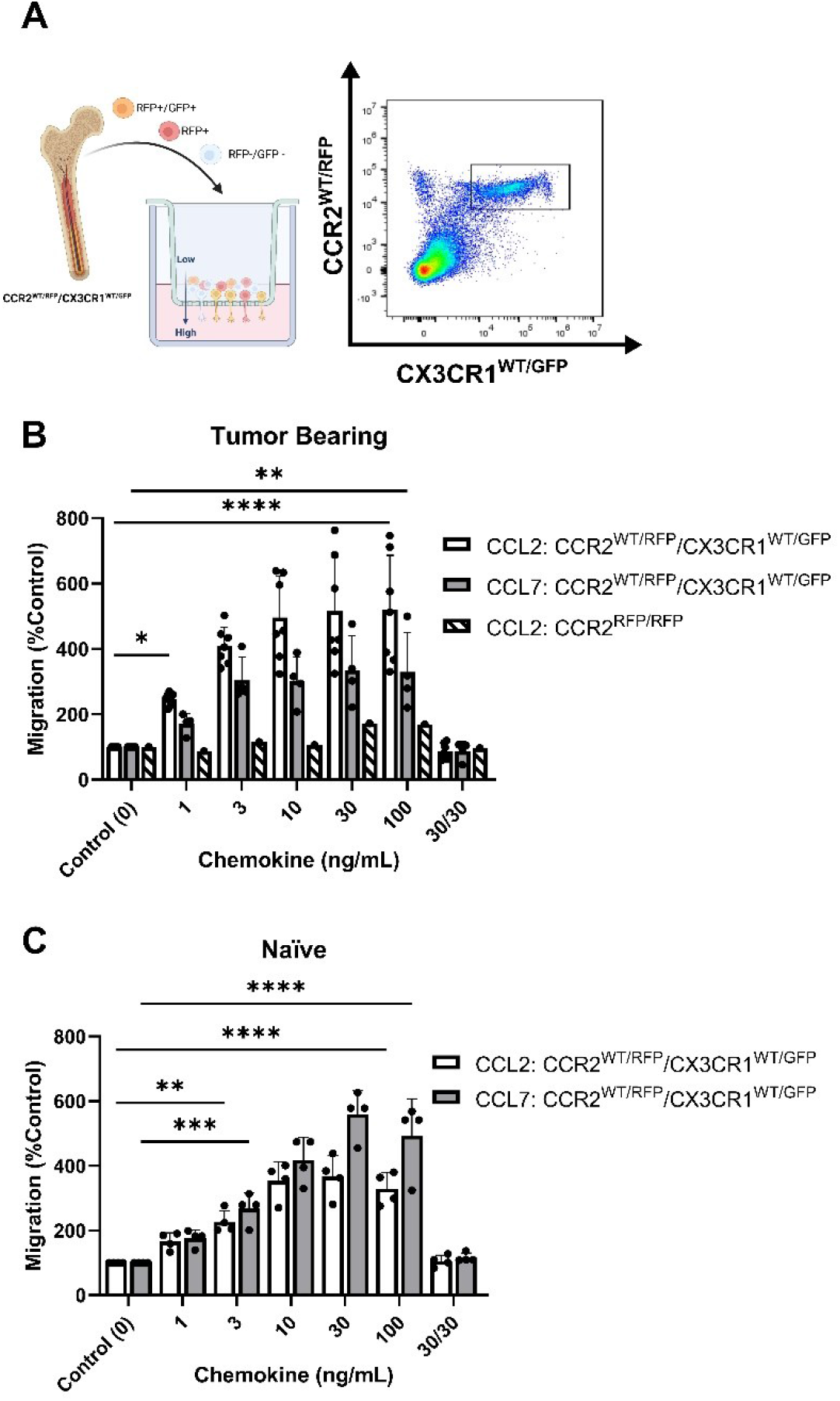
Bone marrow-derived CCR2^+^/CX3CR1^+^ cells migrate to recombinant CCL2 and CCL7 through CCR2. **(A)** Experimental design of transwell migration assays. Graphic (Created with BioRender.com) depicting assay preparation in which whole bone marrow is plated in the top chamber of the transwell migration plate (left). The bottom chamber contains either recombinant chemokine protein or conditioned media. Representative flow plot depicting the population of CCR2^+^/CX3CR1^+^ cells quantified (right). **(B)** Migration to recombinant CCL2 (n=7) and CCL7 (n=4) of CCR2^+^/CX3CR1^+^ cells derived from tumor-bearing animals. Graph also depicts no migration to recombinant CCL2 of bone marrow-derived RFP-expressing cells from Ccr2^RFP/RFP^ animals. **(C)** Migration to recombinant CCL2 and CCL7 of CCR2/CX3CR1-expressing cells derived from naïve animals (n=4). A condition in which chemokine was also plated in the top chamber of the transwell plate (30/30) is included to validate that migration is due to chemotaxis rather than chemokinesis. Two-way ANOVA statistical analysis was conducted (Dunnett’s multiple comparisons test). Differences are compared to the control (0) condition. p-values: 0.0332(*), 0.0021(**), 0.0002(***), <0.0001(****)

Flow cytometry analysis revealed statistically significant migration of CCR2^+^/CX3CR1^+^ cells to recombinant CCL2 and CCL7 (**Figure 4B, C**). Cells derived from naïve animals achieved a maximum percent migration to both ligands at a plating concentration of 30ng/mL (p<0.0001). These cells displayed a higher efficacy for CCL7 relative to CCL2, achieving a mean percent migration of 559% and 366% to CCL7 and CCL2 respectively (**Figure 4C**). Bone marrow-derived cells from tumor-bearing animals 3-week post-implantation also displayed statistically significant migration, achieving maximum migration of 500% and 334% at a plating concentration of 10ng/mL and 30ng/mL for CCL2 and CCL7 respectively (p<0.0001) (**Figure 4B**). Distinct from the naïve condition, cells from the tumor-bearing animal migrate to CCL2 with a slightly higher potency as compared to CCL7 and achieve near-maximum migration to both ligands at as low as 3ng/mL of recombinant protein plated in the bottom chamber.

To establish that the effect shown through these transwell migration assays was due to chemotaxis (directed cell movement in response to a chemokine gradient) as opposed to increased chemokinetic (random cell movement) activity, recombinant chemokine was plated at equal concentrations in both chambers. Disruption of the chemokine gradient, at 30ng/mL concentrations of ligand prevented migration of bone marrow cells derived from either naïve or tumor-bearing animals (**Figure 4B, C**). To determine if the migration was dependent on functional CCR2, bone marrow-derived cells from tumor-bearing CCR2-deficient mice (*Ccr2*^*RFP/RFP*^) were analyzed for CCL2-dependent migration. RFP-expressing cells from *Ccr2*^*RFP/RFP*^ mice did not migrate to any of the CCL2 concentrations tested (**Figure 4B**). This indicates that CCR2-expressing cells migrate to the chemokines CCL2 and CCL7 in a CCR2-dependent mechanism. Since this cell population also expresses CX3CR1, migration of bone marrow-derived cells from tumor-bearing *Ccr2*^*+/RFP*^*/Cx3cr1*^*+/GFP*^ mice to soluble CX3CL1 was assessed. There was no statistically significant migration of CCR2^+^/CX3CR1^+^ cells to CX3CL1 (**Supplemental Figure 5B**). Taken together, these results suggest that bone marrow-derived CCR2^+^/CX3CR1^+^ cells from naïve and tumor-bearing animals migrate to CCL2 and CCL7, in a CCR2-dependent manner.

### KR158B-CCL2 and -CCL7 knockdown gliomas contain equivalent percentages of CCR2^+^/CX3CR1^+^ MDSCs compared to KR158B gliomas

To evaluate whether KR158B tumor cells are active contributors in the recruitment of CCR2^+^/CX3CR1^+^ cells, we tested whether glioma cells produced and secreted CCL2 and CCL7. ELISA analysis of the conditioned media of KR158B cells determined that after 24 hours, glioma cells plated at 500 cells/uL had produced 11.1ng/mL of CCL2 and 1.9ng/mL of CCL7. Analysis of KR158B CCL2 knockdown (KR158B CCL2 KD) and KR158B CCL7 knockdown (KR158B CCL7 KD) cell lines revealed a statistically significant decrease in production of CCL2 and CCL7, measured at 2.7ng/mL and 0.8ng/mL respectively (p<0.0001) (**Figure 5A, B**).

**Figure 5.**
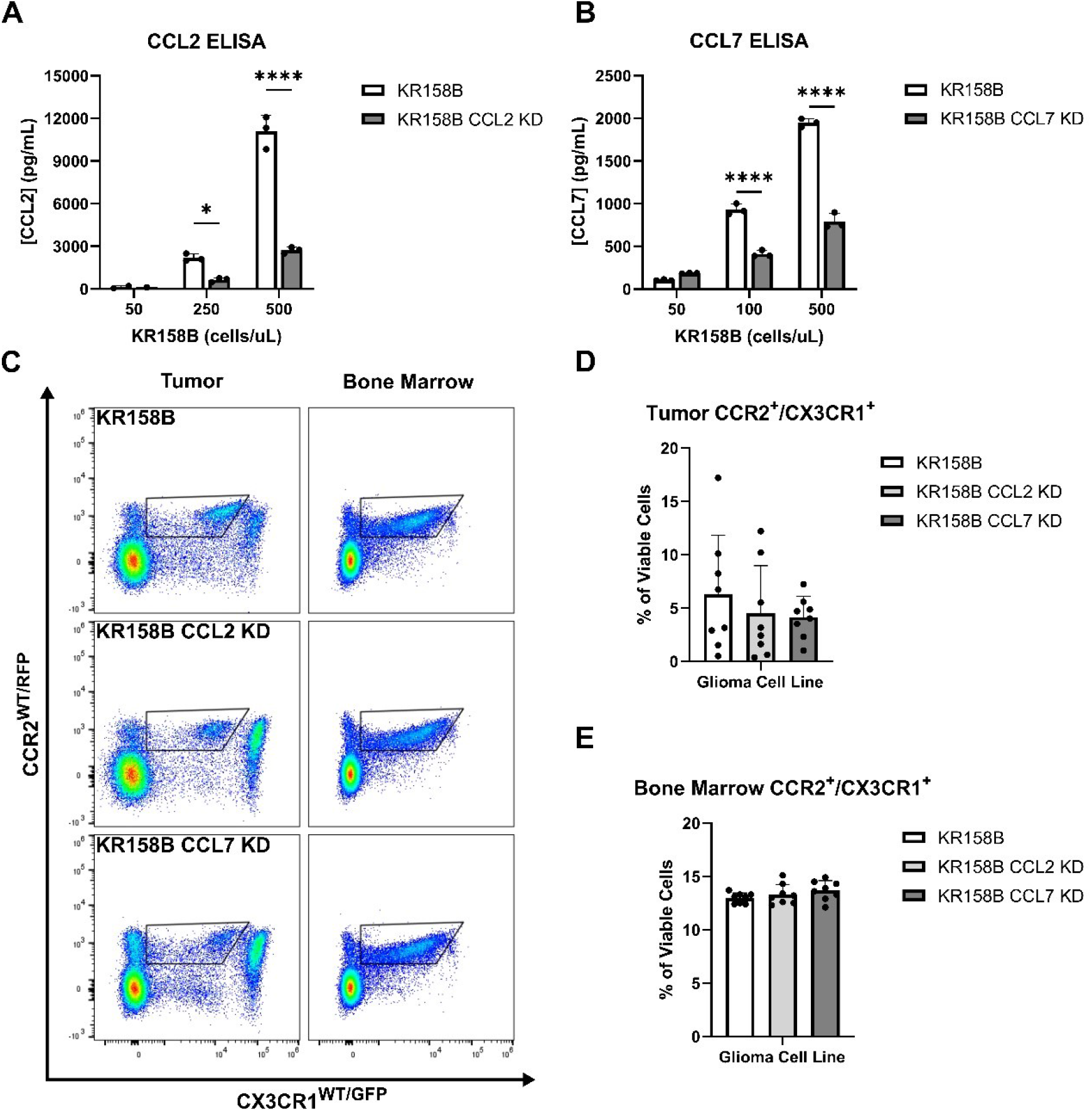
Infiltration of CCR2^+^/CX3CR1^+^ cells into tumors is equivalent in mice implanted with KR158B, KR158B CCL2 KD, or KR158B CCL7 KD tumors. **(A-B)** ELISA results confirming the production of **(A)** CCL2 and **(B)** CCL7 respectively from the KR158B and respective KR158B-chemokine knockdown glioma cell lines. **(C)** Representative flow plots depicting equivalent populations of CCR2/CX3CR1-expressing cells in the chemokine knockdown tumors compared to the KR158B tumor. No significant change in this population was observed in the bone marrow. **(D-E)** Graphs depicting the percent of CCR2/CX3CR1-expressing cells in the tumors and bone marrow of mice implanted with KR158B, KR158B CCL2 KD or KR158B CCL7 KD cell lines (n=8). Two-way ANOVA statistical analysis was conducted (Dunnett’s multiple comparisons test). p-values: 0.0332(*), 0.0021(**), 0.0002(***), <0.0001(****)

KR158B, KR158B CCL2 KD or KR158B CCL7 KD glioma cell lines were implanted in *Ccr2*^*+/RFP*^*/Cx3cr1*^*+/GFP*^ mice, and flow cytometry analysis of tumors and bone marrow was conducted 4.5 weeks post-implantation (**Figure 5C**). We found no significant differences in infiltrating populations of CCR2^+^/CX3CR1^+^ cells, induced by KR158B CCL2 KD and KR158B CCL7 KD glioma cell lines, when compared to KR158B (**Figure 5D**). We also saw no changes in the population of CCR2^+^/CX3CR1^+^ cells in the bone marrow of mice harboring chemokine knockdown gliomas (**Figure 5E**). These results suggest that decreased production of a single chemokine by glioma cells does not impact recruitment of CCR2^+^/CX3CR1^+^ cells to the TME. The lack of effect observed following the implantation of individual CCL2 or CCL7 knockdown gliomas suggested a potential chemokine ligand redundant mechanism utilized by the KR158B cells to recruit CCR2^+^/CX3CR1^+^ cells to the TME.

### CCR2^+^/CX3CR1^+^ cell migration to KR158B conditioned media is inhibited with CCL2 and CCL7 neutralizing antibodies

We next sought to determine whether bone marrow-derived CCR2^+^/CX3CR1^+^ cells migrate to KR158B conditioned media. Utilizing the same transwell migration assay and flow cytometry gating strategy as described above, conditioned media was plated as the chemoattractant, and migration was analyzed. These results indicate that CCR2^+^/CX3CR1^+^ cells migrate significantly to the conditioned media of both the KR158B and KR158B CCL2 KD glioma cell lines. CCR2^+^/CX3CR1^+^ cells migrated to the conditioned media of the KR158B cell line with a maximum percent migration of 266% relative to the migration buffer control condition (p=0.0006) (**Figure 6A**). Consistent with our in vivo results, this cell population migrated similarly to the conditioned media of the KR158B CCL2 KD cell line, achieving a maximum percent migration of 242% relative to the control (p=0.004). Of note, statistically significant migration was only achieved in conditions in which the conditioned media of KR158B or KR158B CCL2 KD cells was derived from 24-hour cultures plated at a concentration of 500cells/uL. A positive control with recombinant CCL2 at 10ng/mL as the chemoattractant confirmed the migratory potential of the cells.

**Figure 6.**
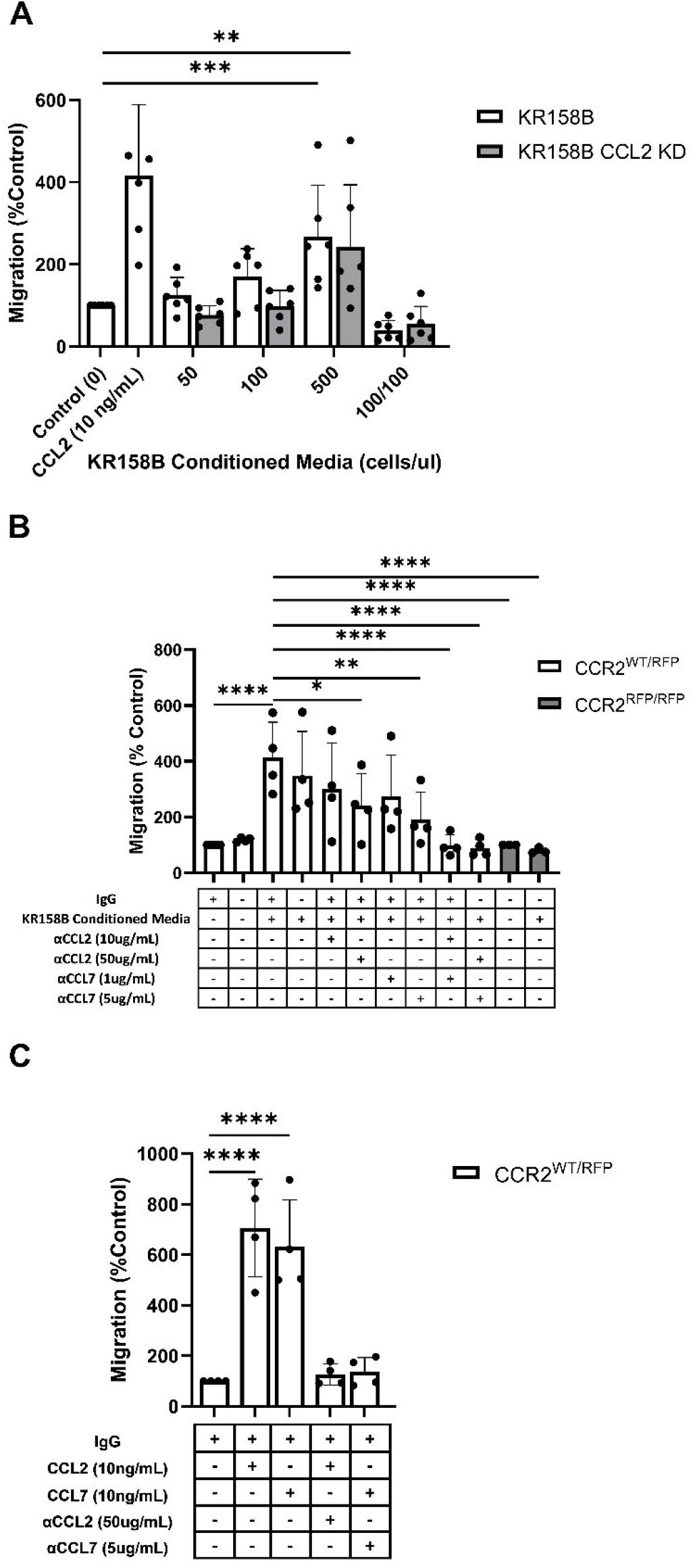
Glioma-derived CCL2 or CCL7 is necessary and sufficient for bone marrow-derived CCR2/CX3CR1 cell migration. **(A)** Graph depicting CCR2^+^/CX3CR1^+^ cells migrating to conditioned media of KR158B (n=6) and KR158B CCL2 knockdown (n=6) cells. A condition in which conditioned media was also plated in the top chamber of the transwell plate (100/100) is included to validate that migration is due to chemotaxis rather than chemokinesis. **(B)** Migration of CCR2^+^/CX3CR1^+^ cells to conditioned media in the presence or absence of chemokine-neutralizing antibodies. Migration is disrupted with the addition of high concentrations of single neutralizing antibody. Migration is completely inhibited with a combination of neutralizing antibodies at either low or high concentrations (n=4). No migration was observed to conditioned media of bone marrow-derived RFP-expressing cells from *Ccr2*^*RFP/RFP*^ animals (n=3). **(C)** Graph depicting that migration to exogenous recombinant CCL2 or CCL7 is inhibited through the addition of high concentrations of respective neutralizing antibody (n=4). Two-way ANOVA statistical analysis was conducted (Dunnett’s multiple comparisons test). Differences are compared to the control condition or between cell lines. p-values: 0.0332(*), 0.0021(**), 0.0002(***), <0.0001(****)

Upon validating CCR2^+^/CX3CR1^+^ cell migration to the conditioned media of KR158B and KR158B CCL2 KD glioma cell lines, we sought to determine whether the migration was exclusively mediated by CCL2 and/or CCL7. To evaluate this question, we plated KR158B cells at 500cells/uL and collected the conditioned media after 24 hours. The impact of anti-CCL2 or anti-CCL7 neutralizing antibodies or a combination of both was evaluated on migration to the KR158B conditioned media. CCR2^+^/CX3CR1^+^ cells migrated significantly to conditioned media containing non-immune IgG with a percent migration of 413% (p<0.0001) (**Figure 6B**). There was no statistically significant difference observed between migration to conditioned media with or without non-immune IgG. In the conditions in which conditioned media was supplemented with a low concentration of neutralizing antibody (10ug/mL α-CCL2 or 1ug/mL α-CCL7), we observed no significant reduction in overall migration. To determine if migration could be inhibited by supplementing with a higher neutralizing antibody concentration, the conditioned media was supplemented with 50ug/mL α-CCL2 or 5ug/mL α-CCL7. When supplementing with 50ug/mL α-CCL2, a significant reduction in percent migration from 413% in the control condition to 240%, a 42% reduction (p=0.016) was evident. Similarly, when supplementing the conditioned media with the high dose of α-CCL7, 5ug/mL, migration was reduced from 413% to 192%, a 54% reduction (p=0.0015).

As CCR2^+^/CX3CR1^+^ cells migrate similarly to CCL2 and CCL7, we then assessed whether there was redundancy between the chemokines that would facilitate cell migration in the event that one ligand was neutralized. To evaluate this, the conditioned media was supplemented with either low combinations (10ug/mL α-CCL2 and 1ug/mL α-CCL7) or high combinations (50ug/mL α-CCL2 and 5ug/mL α-CCL7) of neutralizing antibodies. Interestingly, supplementing with either the low or high combination of neutralizing antibodies resulted in complete inhibition of migration. In the case of the low neutralizing antibody combination, CCR2^+^/CX3CR1^+^ cells achieved 99% migration, whereas cells in the high neutralizing antibody condition reached only 89% migration compared to the migration buffer control normalized to 100% (p<0.0001) (**Figure 6B**).

To confirm that CCR2 was responsible for the migration of CCR2^+^/CX3CR1^+^ cells to KR158B conditioned media, bone marrow cells derived from a *Ccr2*-deficient mouse were utilized. Using conditioned media as the chemoattractant, no statistically significant migration of RFP-expressing cells to conditioned media, as compared to the buffer control, was seen (**Figure 6B**). Taken together, these results show that CCR2^+^/CX3CR1^+^ cells migrate to CCL2 and CCL7 produced by KR158B cells through CCR2, and this migration can be prevented through pharmacologic or genetic disruption of this chemokine-receptor axis.

## Discussion

Glioblastoma is a highly aggressive disease which exhibits a significant immune suppressed tumor microenvironment, lending to its poor prognosis (32–36). Although representing a diverse population in itself, infiltrating myeloid cell populations contribute to the suppressed environment and promote tumor growth (7,8,37–39). Our earlier studies established that CCR2^+^/CX3CR1^+^ myeloid cells, characterized by M-MDSC markers, are present in the bone marrow and infiltrate into multiple murine gliomas (KR158B and 005 GSC). Genetic and pharmacologic inhibition of CCR2 reduced the presence of these cells in the TME, promoted sequestration of these cells in the bone marrow, and unmasked an effect of an immune checkpoint inhibitor to slow glioma progression (26). While these prior studies clearly support targeting CCR2^+^/CX3CR1^+^ cells as a means to treat gliomas, a greater appreciation of the immune suppressive and migratory properties of this CCR2^+^/CX3CR1^+^ cell population is needed. Herein, we extend our published results to better understand the functionality of these cells. The principal findings of this study are 1) glioma-associated CCR2^+^/CX3CR1^+^ myeloid cells are sourced from the bone marrow, 2) CCR2^+^/CX3CR1^+^ cells suppress both CD4^+^ and CD8^+^ T cells, and 3) CCR2^+^/CX3CR1^+^ cells migrate to recombinant and glioma-produced CCL2 and CCL7 in a redundant manner.

Brain tumors, and particularly gliomas, contain mixed populations of myeloid cells (40–44). Our previous report provided a comprehensive analysis of the phenotypic markers expressed by myeloid cells in the glioma microenvironment of both KR158B and 005 GSC intracranial tumors. Glioma-associated myeloid cells can be distinguished by relative CD45 expression, with microglia and peripherally sourced cells expressing mid and high levels of this marker, respectively (26,45). In addition, forward scatter properties also distinguish microglia from peripheral tumor-associated macrophages. We established that bone marrow and glioma-associated CD45^high^, CD11b^+^, Ly6C^hi^/Ly6G^-^ cells co-express CCR2 and CX3CR1. This bone marrow population expands in tumor bearing mice and pharmacological or genetic disruption of CCR2 promotes the sequestration of these cells in the bone marrow (26). Using a chimeric mouse paradigm, we formally established that the CCR2^+^/CX3CR1^+^ population is derived from the bone marrow. These findings suggest an involvement of CCR2, upon stimulation by its ligands, in facilitating the trafficking of CCR2^+^/CX3CR1^+^ cells from the bone marrow to the TME.

CCR2^+^ M-MDSCs represent a prominent infiltrating immune suppressive cell population within murine gliomas (31,46). Their elevated presence has shown to be correlated with negative prognosis and poor response to prospective immunotherapy approaches such as immune-checkpoint inhibitors (9,24). Data reported here establish that CCR2^+^/CX3CR1^+^ M-MDSCs are directly involved in disrupting the proliferation and activated function of both CD4 and CD8-expressing T cells. CCR2^+^/CX3CR1^+^ M-MDSCs suppressed both T cell populations with similar potency. These ex vivo studies are consistent with our prior results where combined PD-1 and CCR2 blockade led to decreased numbers of exhausted CD4^+^ and CD8^+^ T cells and increased IFN-γ expression within the gliomas. Further studies will be necessary to better understand direct and indirect mechanisms whereby CCR2^+^/CX3CR1^+^ M-MDSCs disrupt T cell function and dampen immune responses in the context of glioma. Nonetheless, these data provide further rationale for preventing the infiltration of these immunosuppressive cells into the TME.

CCR2 is a receptor that, among other functions, is primarily implicated in the chemotaxis of cells on which it is expressed (47,48). A common feature amongst many chemokine receptors is the ability to be stimulated by multiple structurally similar ligands. CCR2 is no exception with five known ligands: CCL2, CCL7, CCL8, CCL13 and CCL16 (49). This feature facilitates functional redundancy in that multiple ligands may induce similar downstream cellular effects upon signaling through the same receptor. While CCL2 has previously been reported as the most potent inducer of CCR2^+^ monocyte migration, other CCR2 ligands are likely contributing to migration in a redundant manner to respond to inflammation (50). Brait et al. reported elevated levels of CCL7, in addition to CCL2, in models of ischemia reperfusion (51). These findings suggest that CCL2 and CCL7 may function in a redundant manner to recruit CCR2-expressing cells to sites of inflammation. Additional studies have also reported that the accumulation of myeloid cells within the CNS during inflammation is dependent upon the presence of CCL2 and CCL7. In a CCL2- and CCL7-deficient mouse models, there was a significant reduction in CD45^+^/CD11b^+^/Ly6C^hi^ cells that accumulated in the CNS; the markers that characterize this population coincide with CCR2^+^/CX3CR1^+^ cells in the model utilized here. This suggests that in addition to having functional CCR2, it is also necessary to maintain sufficient levels of the cognate ligands to ultimately induce accumulation of this cell population in the CNS (52). Other chemokine:chemokine receptor systems appear to have redundant roles, determined from studies in murine glioma models, including CCR1 and CCR5 and their shared ligands (53,54). While we report redundant roles for CCL2 and CCL7, there may be spatiotemporal regulation of CCR2-expressing cells by these individual ligands. For instance, one chemokine, i.e., CCL2, might be the prominent driver of CCR2^+^/CX3CR1^+^ cell recruitment to the tumor, while the second, i.e., CCL7, is more important for homing to specific niches within tumor. Although soluble CX3CL1 did not stimulate migration of the MDSCs, a role for membrane attached CX3CL1 in firm adhesion of the cells to endothelium within the tumor vasculature remains a possibility (55,56). Studies aimed toward determining specific CCL2-, CCL7-, and CX3CL1-expressing areas within the tumor would need to be conducted to support these concepts.

In conclusion, we determined that CCR2 and its cognate ligands are prominent regulators of the recruitment of a CCR2^+^/CX3CR1^+^ immune suppressive cell to gliomas. The expression and functional characterization of these chemokine receptors further defines the M-MDSC phenotype. CCL2 and CCL7 are produced, at least in part, by glioma cells and our study indicates that CCL2 and CCL7 function in a redundant manner to induce the migration of CCR2^+^/CX3CR1^+^ M-MDSCs. As such, a more effective approach to limiting this population from gaining access to the TME should involve antagonizing CCR2. However, given that high CCL2 and CCL7 expression is associated with poorer prognosis in GBM patients, consideration of the relative expression of these two chemokines may provide predictive value to a therapeutic strategy targeting this chemokine:chemokine receptor axis.

## Author Contributions

GPT and CJK-designed the study, performed experiments, data analysis, data interpretation, and wrote the manuscript. DL, JSG-performed experiments. GT-performed experiments, data analysis. LPD-designed the study, data interpretation, and wrote the manuscript. DAM-designed the study, data interpretation, and wrote the manuscript. JKH-designed the study, data interpretation, and wrote the manuscript.

## Acknowledgments

We thank Joseph Flores-Toro for providing insight and technical assistance in generating chimeric mice and flow cytometry as well as members of Jeffrey Martens’ lab for access and training on Nikon Confocal Microscope. We gratefully acknowledge the University of Florida Interdisciplinary Center for Biotechnology Research Flow Cytometry Core and University of Florida Animal Care Service.

