## Supplemental Material for "Glioma-derived CCL2 and CCL7 mediate migration of immune suppressive CCR2^+^ myeloid cells into the tumor microenvironment in a redundant manner"

### Supplementary Figures

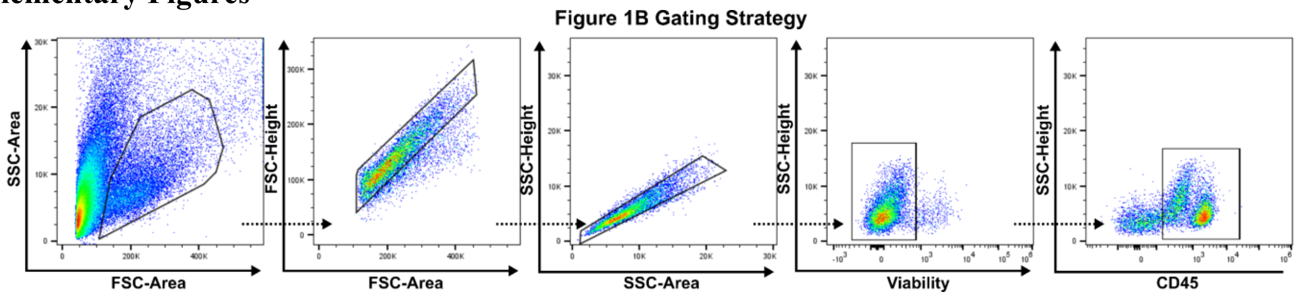

**Supplementary Figure 1.** Representative flow cytometry gating strategy for Figure 1B analysis of tumor infiltrating CCR2$^{+}$/CX3CR1$^{+}$ cells in control and chimeric mice. Identical gating strategy was applied to all other conditions.

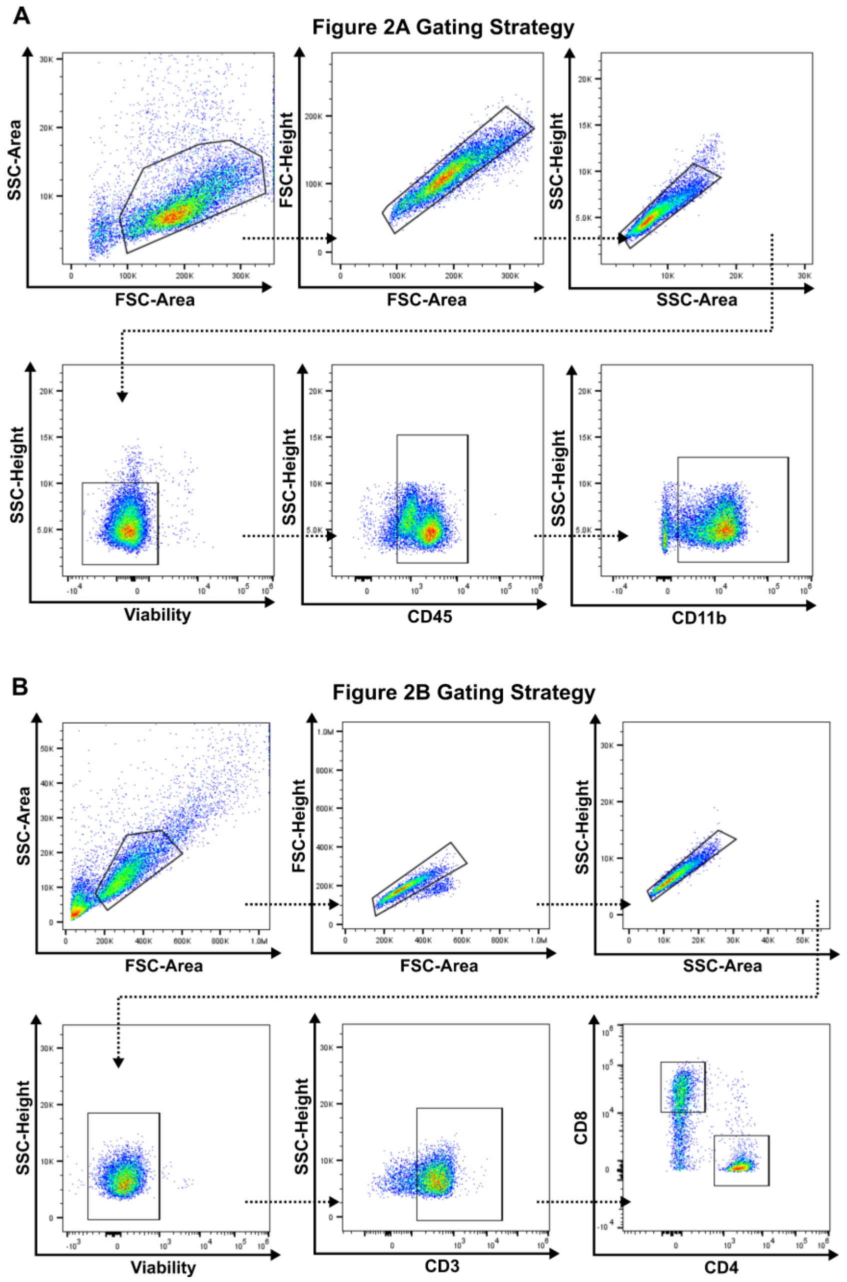

**Supplementary Figure 2. (A)** Representative flow cytometry gating strategy for Figure 2A post magnetically activated cell sorting. Identical gating strategy was applied to all other conditions in that panel. **(B)** Representative flow cytometry gating strategy for Figure 2B T cell suppression assay. CD4 and CD8 T cell proliferation were analyzed simultaneously. Identical gating strategy was applied to all other conditions in that panel.

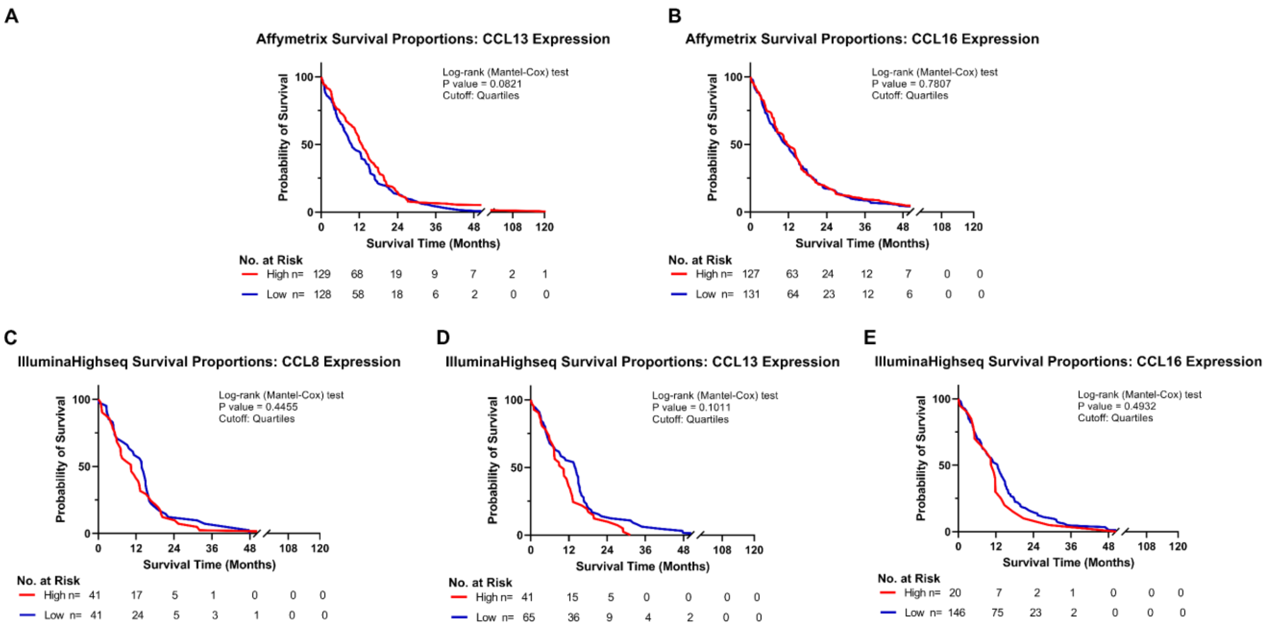

**Supplementary Figure 3. Additional human CCR2 chemokine ligands do not confer negative prognosis for patients with glioblastoma. (A-B)** Kaplan-Meyer survival curves of GBM patients based on Affymetrix gene expression profiles of **(A)** CCL13 **(B)** CCL16 **(C-E)** Kaplan-Meyer survival curves of GBM patients based on Illumina Highseq expression profiles of **(C)** CCL8 **(D)** CCL13 **(E)** CCL16 mined from TCGA database. High and low cohorts are stratified as top and bottom quartiles, respectively. Number at risk indicates surviving patients in each cohort at the respective timepoints of analysis. Log-rank (Mantel-Cox) test was conducted on high vs low expressing cohorts.

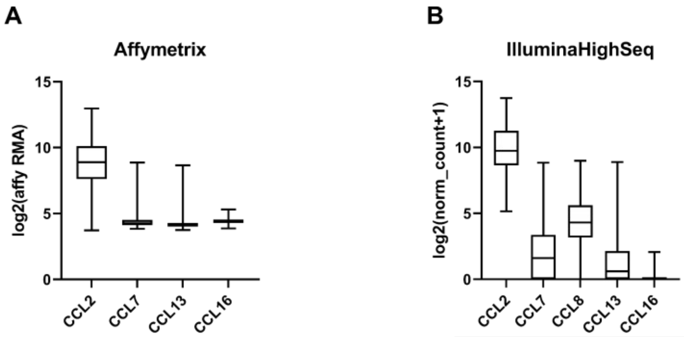

| | CCL2 | CCL7 | CCL13 | CCL16 |
|---|---|---|---|---|
| Number of values | 518 | 518 | 518 | 518 |
| Minimum | 3.722 | 3.842 | 3.760 | 3.869 |
| 25% Percentile | 7.622 | 4.116 | 4.014 | 4.293 |
| Median | 8.893 | 4.259 | 4.151 | 4.420 |
| 75% Percentile | 10.12 | 4.523 | 4.289 | 4.545 |
| Maximum | 12.97 | 8.865 | 8.654 | 5.310 |
| Range | 9.248 | 5.023 | 4.894 | 1.441 |

| | CCL2 | CCL7 | CCL8 | CCL13 | CCL16 |
|---|---|---|---|---|---|
| Number of values | 166 | 166 | 166 | 166 | 166 |
| Minimum | 5.154 | 0.000 | 0.000 | 0.000 | 0.000 |
| 25% Percentile | 8.646 | 0.000 | 3.172 | 0.000 | 0.000 |
| Median | 9.745 | 1.602 | 4.309 | 0.6093 | 0.000 |
| 75% Percentile | 11.27 | 3.371 | 5.615 | 2.138 | 0.000 |
| Maximum | 13.74 | 8.844 | 8.993 | 8.898 | 2.069 |
| Range | 8.584 | 8.844 | 8.993 | 8.898 | 2.069 |

**Supplementary Figure 4.** Descriptive statistics of TCGA GBM survival analysis for CCR2 chemokine ligands CCL2, CCL7, CCL8, CCL13, and CCL16. **(A)** Affymetrix gene expression cohort **(B)** Illumina Highseq gene expression cohort. Number of patients analyzed, median, range, and quartiles are displayed. Quartiles were used to stratify high and low populations in survival analysis.

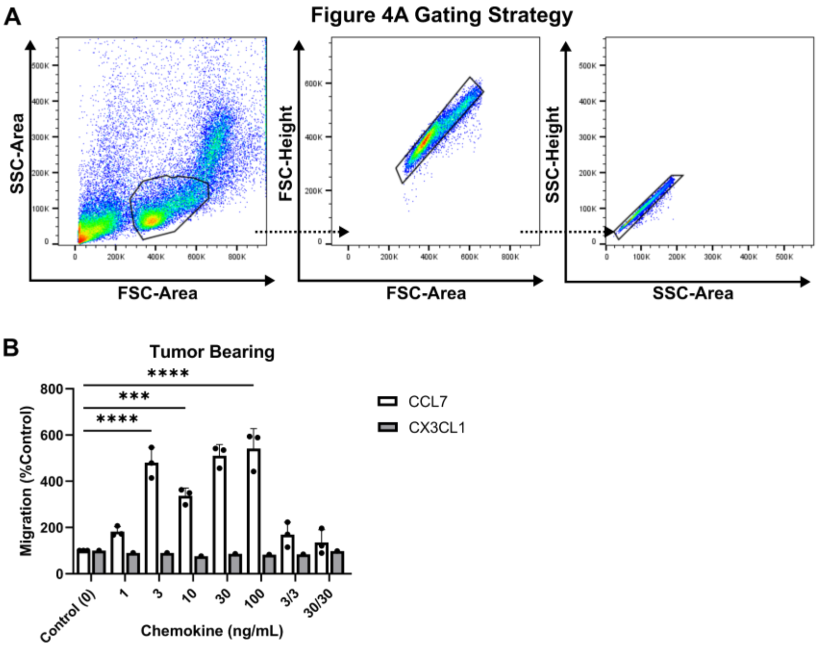

**Supplementary Figure 5. CCR2$^{+}$/CX3CR1$^{+}$ cells do not migrate to soluble CX3CL1. (A)** Representative flow cytometry gating strategy for Figure 4A in vitro migration experiments. Identical gating strategy was applied to all other conditions in that figure. **(B)** Migration to recombinant CCL7 (n=3) and recombinant soluble CX3CL1(n=1) of CCR2/CX3CR1 expressing cells derived from glioma bearing animals. Graph depicts no migration to recombinant CX3CL1. Two-way ANOVA statistics was conducted (Dunnett's multiple comparisons test). Differences are compared to the control (0) condition. p-values: 0.0332(*), 0.0021(**), 0.0002(***), <0.0001(****)

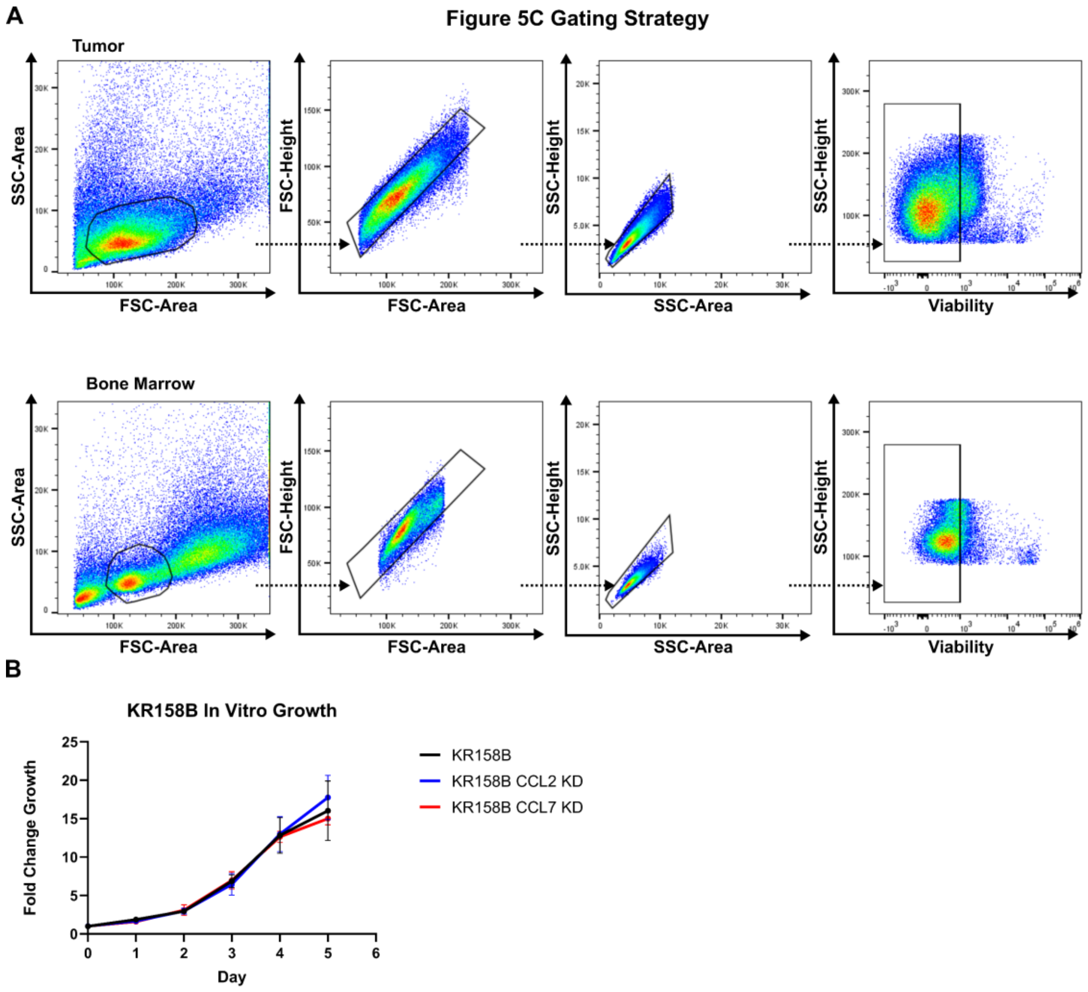

**Supplementary Figure 6. Glioma KR158B CCL2- and CCL7- knockdowns display similar growth in vitro. (A)** Representative flow cytometry gating strategy for Figure 5C in vivo migration of CCR2$^{+}$/CX3CR1$^{+}$ cells: tumor (top) and bone marrow (bottom). Identical gating strategy was applied to all other conditions in that panel. **(B)** In vitro growth analysis of KR158B, KR158B CCL2 KD, and KR158B CCL7 KD glioma cell lines. CyQUANT Cell Proliferation dye was used to assess growth. Data is displayed as fold change growth from day 0 to day 5.

Flow Cytometry

| Protein Marker | Fluorophore | Company | Catalog # | Dilution Factor |
|---|---|---|---|---|
| CellTrace | FarRed | Invitrogen | C34564 | 1ul/1 x $10^6$ cells |
| CD11b | BV421 | Biolegend | 101251 | 1:100 |
| CD3 | BV510 | Biolegend | 100234 | 1:100 |
| CD4 | FITC | Biolegend | 100510 | 1:100 |
| CD45 | Alexa700 | Biolegend | 103128 | 1:200 |
| CD8 | PE | Biolegend | 100708 | 1:100 |
| LY6G | PerCP | Biolegend | 127654 | 1:100 |
| LY6C | BV785 | Biolegend | 128041 | 1:100 |
| Viability | Pacific Blue | Invitrogen | L34963 | 1µl/mL |

Biolegend, San Diego CA; Invitrogen, Carlsbad, CA
